# Toward CT-based Tractography: Presurgical White Matter Tract Mapping in Intracerebral Hemorrhage

**DOI:** 10.64898/2026.05.18.724202

**Authors:** Guanlin Huang, Guoqiang Xie, Yijie Li, Qun Wang, Shun Yao, Yiheng Tan, Ron Kikinis, Alexandra J Golby, Lauren J O’Donnell, Fan Zhang

## Abstract

Presurgical mapping of key white matter (WM) fiber tracts is crucial for intracerebral hemorrhage (ICH) surgery, but it currently relies on tractography from diffusion MRI (dMRI), which has limited applicability in urgent or resource-constrained settings due to long scan times and limited MRI availability. To bridge this gap, we developed a deep learning approach designed to reconstruct critical fiber tracts directly from routinely acquired CT scans, focusing on the corticospinal tract (CST) due to its high clinical relevance. By training a novel network on a curated dataset of 150 paired CT-dMRI scans (101 ICH patients and 49 healthy subjects), we enabled the direct mapping of the CST from CT images alone, bypassing the traditional requirement for dMRI. Our results demonstrate that the CT-derived tracts achieve high anatomical plausibility, with neurosurgeon expert assessments yielding a Likert score of 3.64. Furthermore, the clinical relevance of these reconstructions was validated by a significant correlation between CT-derived tract integrity and patient motor scores (r = 0.726, p = 3.731× 10^−7^). These findings suggest that in complex clinical scenarios, particularly where dMRI signal quality is compromised by lesion-induced distortion, this CT-based mapping may serve as a useful anatomical reference. Overall, by enabling CT-based WM mapping, our method potentially offers a practical route to gain the key advantages of tractography in resource-limited or time-critical settings

## 1. Introduction

Intracerebral hemorrhage (ICH) stroke is a critical medical emergency characterized by high mortality, and its rapid treatment is essential to halt ongoing bleeding, mitigate neuronal damage, and significantly improve clinical outcomes. Accurate identification and preservation of critical white matter (WM) fiber tracts are paramount in the surgical management of ICH ^1–4^. In particular, the corticospinal tract (CST) has high clinical relevance in ICH, where its integrity is critical for the preservation of patient motor function ^5,6^. Utilizing advanced diffusion MRI (dMRI) tractography allows for the mapping of eloquent tracts in relation to the hematoma. In ICH, tractography is a valuable tool to assist presurgical planning and surgical prognosis ^7^. For example, Labib et al. conducted a multicentric randomized prospective trial, which demonstrated that tractography can identify eloquent white matter tracts adjacent to the clot and subsequently guide the planning of the safest surgical approach ^8^. Jang et al. and Kumar et al. demonstrated that tractography helps monitor damage and recovery in eloquent white matter tracts during the subacute phase of hemorrhage, providing insights into long-term functional prognosis ^9^. However, the clinical utility of dMRI in ICH is often limited by long acquisition times and limited scanner availability, particularly in urgent cases and/or under-resourced healthcare environments without MRI scanners. Conversely, computed tomography (CT) is the frontline imaging modality for ICH in routine clinical practice due to its speed and ubiquity, but it offers no inherent fiber tracking capabilities. This disparity underscores a critical unmet need: the development of computational methods to accurately infer the location of critical white matter pathways from routinely acquired CT scans.

Tractography is an advanced neuroimaging technique that is currently the only method for *in vivo* mapping of WM fiber tracts in the living human brain ^10^. Technically, tractography is a computational process that reconstructs the 3D trajectories of WM fiber pathways by tracing the water diffusion directions in axons. Traditional methods use dMRI data to estimate fiber orientations through tissue modeling using techniques such as the classical diffusion tensor model, as well as more advanced approaches like Constrained Spherical Deconvolution (CSD), multi-fiber models, and global tractography ^11^. Recent advances have shifted toward deep learning methods ^12^. For instance, TractSeg ^13,14^ is one of the most popularly used deep learning tractography methods and has proven effective for surgical WM mapping ^15–17^. Rather than using model-fitting approaches, TractSeg employs neural networks to estimate the so-called tract orientation mapping (TOM), which provides a tract-specific fiber orientation representation to directly generate streamline tractograms. While such deep learning methods have improved tractography by enabling fast and accurate fiber tract reconstruction, their reliance on dMRI data remains a major limitation in ICH, where dMRI acquisition times are too long for urgent clinical decision making, and MRI scanners may be unavailable or under extreme use pressure.

Despite its utility, dMRI-based tractography also faces inherent limitations in the context of spontaneous ICH. During the pathological evolution of ICH, perihematomal edema (PHE) serves as a critical driver of secondary brain injury ^18^. The accumulation of interstitial fluid within edematous regions results in a high concentration of isotropic water, which significantly dilutes the anisotropic signals reflecting underlying neural architecture. This phenomenon leads to a precipitous decline in Fractional Anisotropy (FA)—a primary metric of tissue integrity— which conventional tracking algorithms often misinterpret as a white matter boundary, resulting in premature termination of the reconstruction ^19,20^. Furthermore, the mass effect exerted by the hematoma and its surrounding extensive edema frequently causes mechanical displacement or structural disruption of white matter tracts ^21^. Consequently, integrating complementary imaging modalities into the tractography framework is beneficial for achieving more robust anatomical reconstructions.

Recent work increasingly moves beyond traditional dMRI-based tractography, emphasizing the use of more widely available neuroimaging data. The overall idea is to learn mappings between fiber tract information captured in dMRI and signals from alternative imaging modalities, allowing tract reconstruction in the absence of dMRI. For example, multiple studies proposed to leverage anatomical features from T1-weighted MRI ^22,23^ or FLAIR MRI ^24^ to reconstruct WM tracts independently of dMRI data. These advances demonstrate the potential to redefine tractography workflows, yet challenges persist for our interest in reconstructing WM tracts from standard CT scans in ICH patients, due to several fundamental challenges: (1) the inherent low soft-tissue contrast of CT, which provides inadequate differentiation of WM pathways; (2) significant obfuscation of native neuroanatomy in ICH patients from the acute hemorrhage’s mass effect, surrounding edema, and hyperdense signal; and (3) the limited availability of paired CT-dMRI datasets required for training deep learning models. More recently, Murray et al. ^25^ proposed a method to predict the CST from CT scans in ICH patients, providing important evidence that CT-based tractography is feasible and holds significant clinical potential. This work lays a valuable foundation for subsequent research in this direction. However, their method focuses on predicting the presence of white matter tracts as probabilistic maps, rather than reconstructing the detailed 3D fiber trajectories.

In this work, we propose a novel tractography method to reconstruct WM fiber tracts directly from CT scans in ICH patients, eliminating the need for dMRI and its associated challenges of prolonged acquisition times and limited scanner availability. Given the close association of CST with motor function ^5,6^, which has high clinical relevance in ICH, we focus on the CST. Our study has three major contributions. First, we acquire and curate a large, prospective dataset (n=150) that includes paired CT and dMRI scans from not only ICH patients but also healthy subjects (HSs). Second, we design a novel deep learning network that leverages TOM images designed in TractSeg as an intermediate tract representation and performs CT-based tract-specific tractography by learning the mapping from CT to TOM. Third, we demonstrate robust tract identification in ICH CT scans, supported by expert assessment and clinical relevance analyses, with improved performance relative to dMRI-based methods when mass effect is substantial.

## 2. Methods

### 2.1. Study Participants and Image Acquisitions

A total of 150 participants, including 101 ICHs and 49 HSs, were recruited at the department of neurosurgery in Nuclear Industry 215 Hospital of Shaanxi Province, China. The inclusion criteria of this study were as follows: (1) first-ever acute hemorrhagic stroke (for patient only); (2) no psychiatric diseases, conscious disorders, or cognitive deficits; (3) no contradictions for MRI or CT examinations. The institutional review board of the hospital approved the study. Written informed consent was obtained in compliance with the hospital’s ethics policy.

For each participant, paired CT and dMRI scans were prospectively acquired. For the ICH patients, the CT and dMRI data were acquired within 15 days of symptom onset, on average. The two scans were performed within a narrow timeframe of 2 days. Specifically, the hematomas in ICH patients were primarily located in the basal ganglia, and the average hematoma volume was 38 ± 22.8 mL (range: 4–93 mL). For HSs, the CT and dMRI data were obtained on the same day. The acquisition parameters were as follows. The CT data were scanned on a Philips MX 16-slice CT scanner, with voxel size=0.5×0.5×6 mm^3^ for ICH patients and 0.5×0.5×1 mm^3^ for HSs. The dMRI data were acquired on a GE Discovery MR750 scanner, with TE/TR=80.8/8750 ms, 2×2×2 mm^3^ resolution, 1 baseline image at b=0 s/mm^2^ and 30 gradient directions at b=1000 s/mm^2^.

For each ICH patient, limb motor function was assessed (by neurosurgeon GX) at the time of CT imaging using Lovett muscle strength testing ^26^, based on the ability of a muscle to contract and resist gravity/resistance (0: no muscle activation; 5: normal).

### 2.2. Data Preprocessing

The dMRI data underwent standard preprocessing via an established pipeline (https://github.com/pnlbwh/pnlpipe), including brain extraction and eddy current correction. The CT scans were preprocessed using 3D Slicer ^27^ with brain extraction. The dMRI and CT data were co-registered, followed by a rigid registration into the standard MNI space, using ANTs ^28^. Following registration, all imaging data were resampled to a consistent spatial resolution to ensure anatomical alignment and cross-modal voxel-wise correspondence

### 2.3. Ground-Truth Tract Generation

The ground truth TOM images were derived from the processed dMRI data. This was done by computing fiber orientation distribution (FOD) peaks using constrained spherical deconvolution (CSD) implemented in MRtrix ^29^. Subsequently, the FOD peaks were used as input for TractSeg ^13,14^ to generate TOMs of CST for each subject. A TOM captures the local fiber orientation of a tract, with each voxel representing a single 3D vector (or peak) that denotes the mean orientation of the streamlines passing through it. From the computed TOM, we performed streamline tracking to obtain the ground truth CST fiber pathway using TractSeg.

### 2.4. Deep Model for Tract-Specific Tractography on CT

Figure 1 provides a method overview. A deep model is learned on a training dataset with paired CT and dMRI scans, where the dMRI-derived TOMs serve as the ground truth. The trained model is then applied to new CT scans to predict TOMs, enabling subsequent fiber tracking to reconstruct the CST.

**Figure 1.**
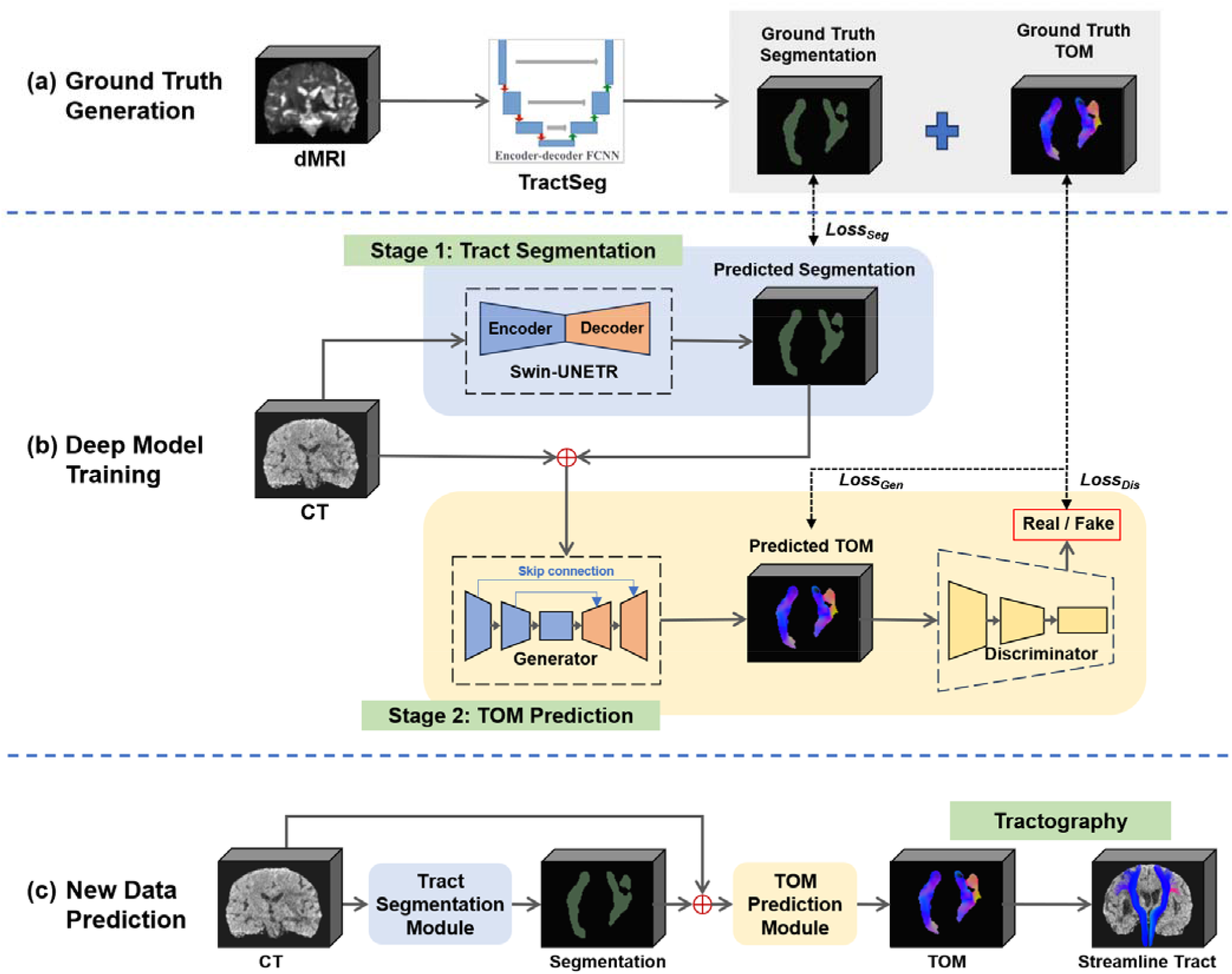
Schematic of the proposed CT-based tractography pipeline. **(a)** Ground Truth Generation: Tract segmentation and Tract Orientation Mapping (TOM) images are derived from diffusion MRI (dMRI) to serve as model training targets. **(b)** Deep Model Training: A two-stage deep model learns to map input CT data to the ground truth by first computing a tract segmentation, followed by the tract TOM prediction. **(c)** New Data Prediction: For a new CT scan, the trained model predicts a tract TOM image, which is then used for streamline tractography to reconstruct fiber pathways.

We design a novel two-stage network architecture that first localizes the tract of interest via segmentation and then uses this segmentation as a spatial prior to guide the prediction of the TOM, as follows. First, we generate a segmentation of the WM tract of interest from the CT. This is done by training a Swin-UNETR ^30^ using the CT images as input and the binarized tract mask as segmentation ground truth. Then, we employ a conditional Generative Adversarial Network (cGAN) to predict the TOM images. To guide the generation towards specific anatomical regions, we use the pre-trained Swin-UNETR model as a backbone network to produce the tract segmentation as a spatial prior. The generated tract segmentation is channel-concatenated with the CT scan to provide spatial guidance to a 3D PatchGAN ^31^ for TOM prediction. This conditioning directs the model’s focus to the relevant tract regions, thereby enhancing the quality of the generated output.

#### 2.4.1. Tract Segmentation Module

The tract segmentation module is built based on the SwinUNETR architecture, which integrates Swin Transformer blocks within a U-Net style encoder-decoder framework. The network takes 80×80×80 CT patches as input and processes them through Transformer-based encoder stages, which capture long-range contextual dependencies via shifted window self-attention. The encoder gradually expands features through four layers of Swin Transformer blocks, while skip connections transmit multi-scale spatial information to the corresponding decoder layers. Finally, a linear projection layer followed by a sigmoid activation generates the tract segmentation.

#### 2.4.2. TOM Prediction Module

The TOM prediction module is built on the conditional GAN architecture. The generator utilizes an attention UNET backbone, introducing attention gates at each unsampling level of the decoder to effectively suppress noise regions in the input image and focus on the reconstruction of anatomical structures. At the input, a dual-channel configuration is employed, concatenating the original CT image with the anatomical segmentation, serving as a spatial prior to guide the generation process. Rather than processing the entire volume, the discriminator incorporates the design of the PatchDiscriminator, which uses three consecutive convolutional layers to assess the authenticity of local patches of the generated volume. This patch-based evaluation method is more capable of capturing high-frequency details and textures compared to traditional global discriminators, ensuring that the generated images are consistent with real data at a microscopic structural level.

#### 2.4.3. Loss Functions

For training segmentation module, a Dice loss function is adopted, as:

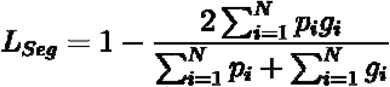

where *N* is the total number of voxels, *p* is the predicted tract segmentation and *g* is the ground truth segmentation obtained from dMRI.

For training TOM prediction module, the generator loss function L_*Gen*_ combines the L_1_ loss and adversarial loss to ensure that the generated TOM has higher anatomical accuracy, while the discriminator loss function L_*Dis*_ employs the least squares adversarial loss to make the generated TOM closer to the distribution of the ground truth, as:

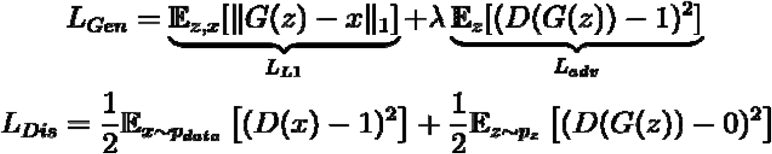

where *x* is the ground truth TOM, *G(z)* is the generated TOM, λ is the weight coefficient of adversarial loss.

### 2.5. Implementation and Parameter Settings

We implement the model using PyTorch 2.5 on an NVIDIA RTX 3090 GPU with 24 GB of memory. The network is trained for 200 epochs with the AdamW optimizer (β□ = 0.9, β □ = 0.999). The learning rates are set to 1×10□ □ for the generator and 5×10□ □ for the discriminator. Training is performed on 3D patches of size 80×80×80 voxels, using a batch size of 3 with a queue-based data loading mechanism (queue capacity: 50 volumes; 10 patches sampled per volume). During inference, an overlapping grid sampling strategy with a 20-voxel overlap is applied to ensure seamless volume reconstruction.

## 3. Statistical Analyses

First, we evaluate network design using three comparative experiments: (1) an ablation of segmentation-guided conditioning (with or without it), (2) an assessment of training data composition (disease-only vs. disease+healthy), and (3) comparison with baseline deep-learning methods (Fully Convolutional Neural Network (FCNN) ^14^ and Denoising Diffusion Probabilistic Model (DDPM) ^32^). Predictions are evaluated against ground-truth TOMs and tracts derived from dMRI in ICH (n=20) and healthy (n=10) test data. To rigorously assess model performance under pathological distortion, evaluations are conducted on both the hemispheres affected by ICH and the contralateral unaffected hemispheres. Three metrics are used: *Mean Squared Error (MSE)* measures voxel-wise differences in TOM (lower is better); *Dice Similarity Coefficient (DSC)* assesses volumetric overlap between tracts (higher is better) ^33^; and *Tract Distance (TD)* quantifies streamline geometric agreement between tracts (lower is better) ^34^. In the ablation and training-data experiments, methods are compared using paired t-tests with false discovery rate (FDR) correction across metrics. For baseline comparisons, a one-way repeated-measures ANOVA is performed, followed by post-hoc paired t-tests with FDR correction across metrics.

Second, we perform an expert assessment to evaluate the anatomical plausibility and clinical utility of predicted tracts in the ICH-affected hemisphere. Three independent practicing neurosurgeons (GX, SY, QW) assess tractography by rating their agreement (5-point Likert scale ^35^) with two statements: (1) the tract passes through expected brain regions, and (2) hematoma-induced tract displacement or splitting is accurately represented. For comparison, the same assessment is conducted for dMRI-based tracts generated with TractSeg. To ensure blinding, CT- and dMRI-derived tractography results are randomly shuffled. Corresponding CT, b0, FA, and MD images are provided to support evaluation. Inter-rater consistency is assessed using the Intraclass Correlation Coefficient (ICC) ^36^.

Third, we assess clinical relevance of the predicted CSTs by correlating their tract integrity with motor function in ICH patients, testing the established link between WM damage and neurobehavioral dysfunction ^1,3^. For each test patient (n=20), left and right CST volumes are extracted as markers of tract integrity and correlated with contralateral Lovett muscle strength scale scores using Pearson’s correlation. By pooling data from both the affected and unaffected sides, we ensured a sufficient variance in tract integrity to robustly test the established link between white matter damage and motor dysfunction. Correlation coefficients (*r*) and p-values are reported.

## 4. Results

### 4.1. Dataset Characteristics

Table 1 gives the demographics of the recruited participants. The 150 participants are randomly split into training (n=112; 76 ICH, 36 HS), validation (n=8; 5 ICH, 3 HS), and testing (n=30; 20 ICH, 10 HS) sets, following a 75%/5%/20% split. The dataset is stratified to ensure that hematoma locations are proportionally represented across the training, validation, and testing sets, maintaining a balanced distribution for model evaluation.

**Table 1:**
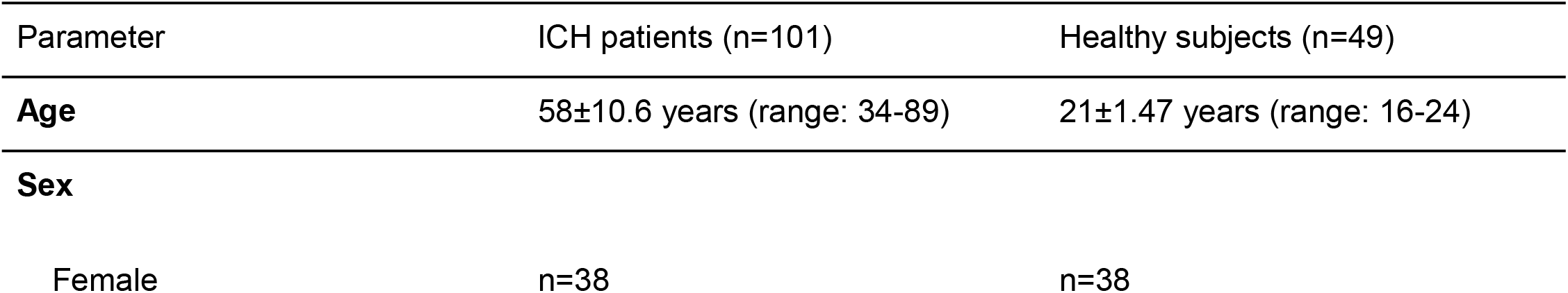
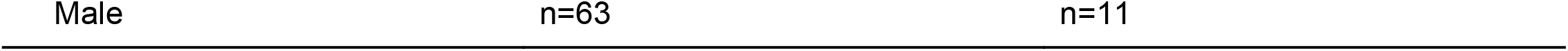
Participant Demographics.

### 4.2. Evaluation of the deep network

To ensure a rigorous and head-to-head comparison, both CT-based and dMRI-based tractography utilize identical seed regions and tracking parameters. Fiber reconstruction is performed using the Fiber Assignment by Continuous Tracking (FACT) algorithm, with a consistent step size and termination criteria (e.g., angular threshold and FA limit) applied across all experimental conditions.

Table 2 summarizes results from the three comparative experiments in both affected and unaffected hemispheres. First, the ablation study shows that segmentation-guided conditioning significantly improves CST prediction, increasing DSC from 0.63 to 0.81 (in unaffected hemispheres), from 0.56 to 0.70 (in affected hemispheres), reducing TOM MSE from 0.133 to 0.084 (in unaffected hemispheres), from 0.152 to 0.105 (in affected hemispheres), and decreasing TD from 6.16mm to 4.56mm (in unaffected hemispheres), from 6.70 mm to 5.61 mm (in affected hemispheres) (all p<0.001, paired t-test, FDR-corrected). Second, the analysis of training data composition demonstrates that incorporating both ICH and HS data significantly improves CST prediction compared to using ICH data alone (all p<0.001, paired t-test, FDR-corrected). Third, baseline comparisons show that our method significantly outperforms FCNN and DDPM, achieving the highest DSC and lowest TOM MSE and TD across all metrics (p<0.001, ANOVA with post-hoc pairwise t-tests).

**Table 2:**
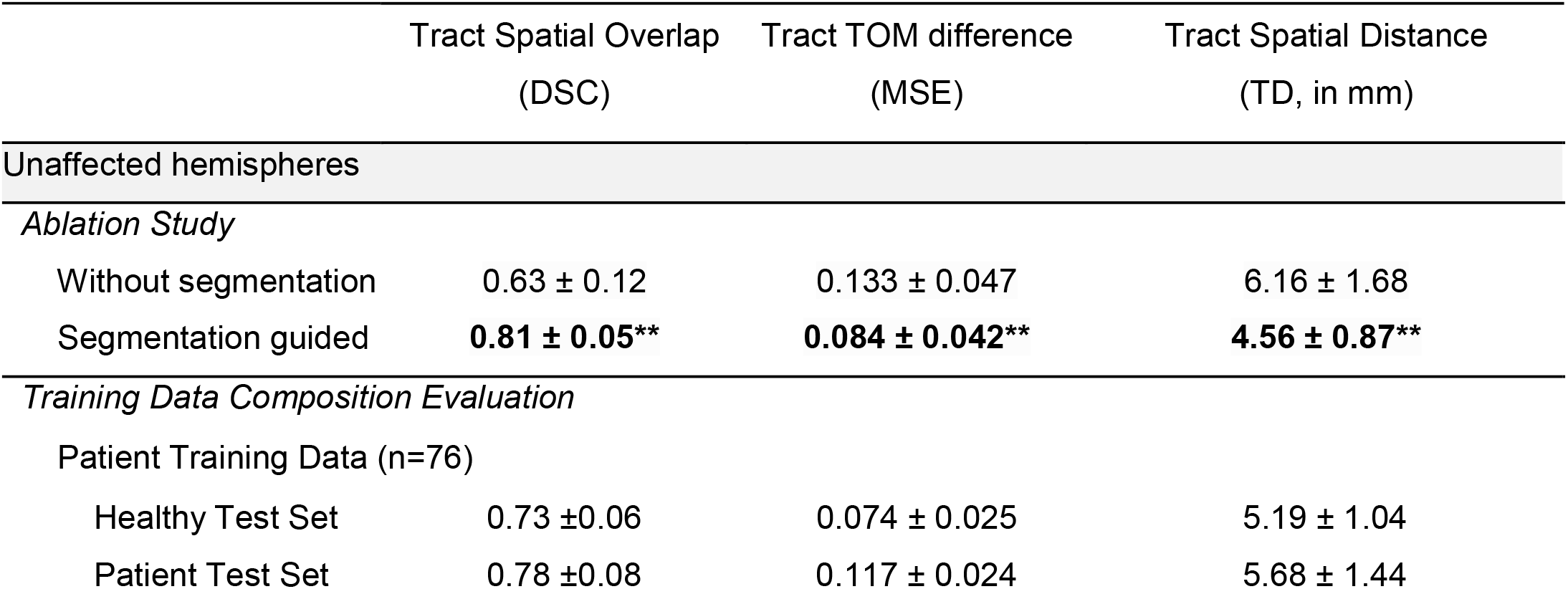
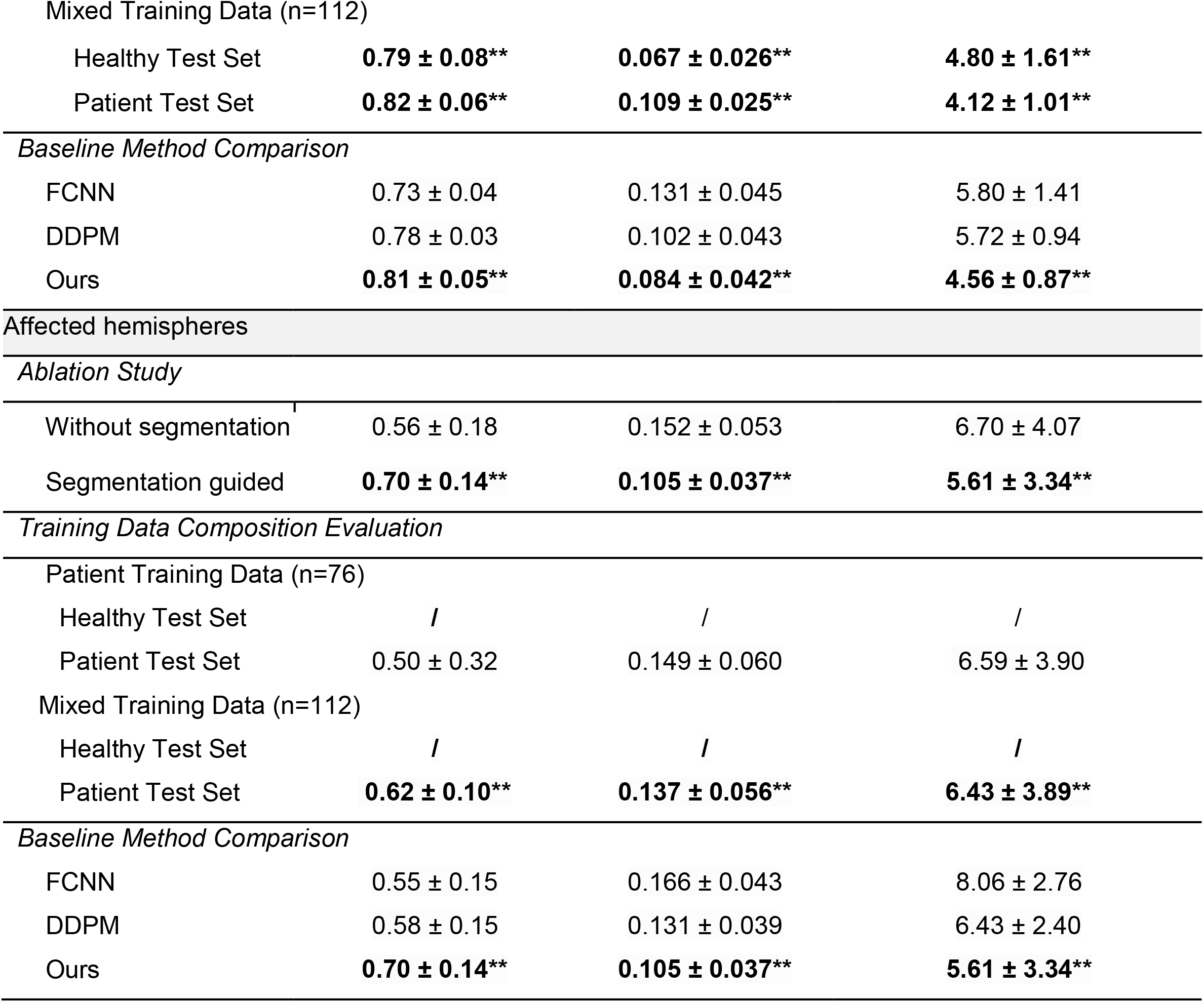
Results of the three comparative experiments.

Figure 2 presents a visual comparison of predicted TOMs and streamline tracts generated by FCNN, DDPM, and our method in representative HS and ICH patients. DDPM exhibits conspicuous stitching artifacts, while FCNN produces overly smooth outputs that miss structural details. In contrast, our method generates TOMs that most closely resemble the dMRI-derived ground truth, resulting in streamline pathways that more accurately represent the underlying anatomical structure.

**Figure 2.**
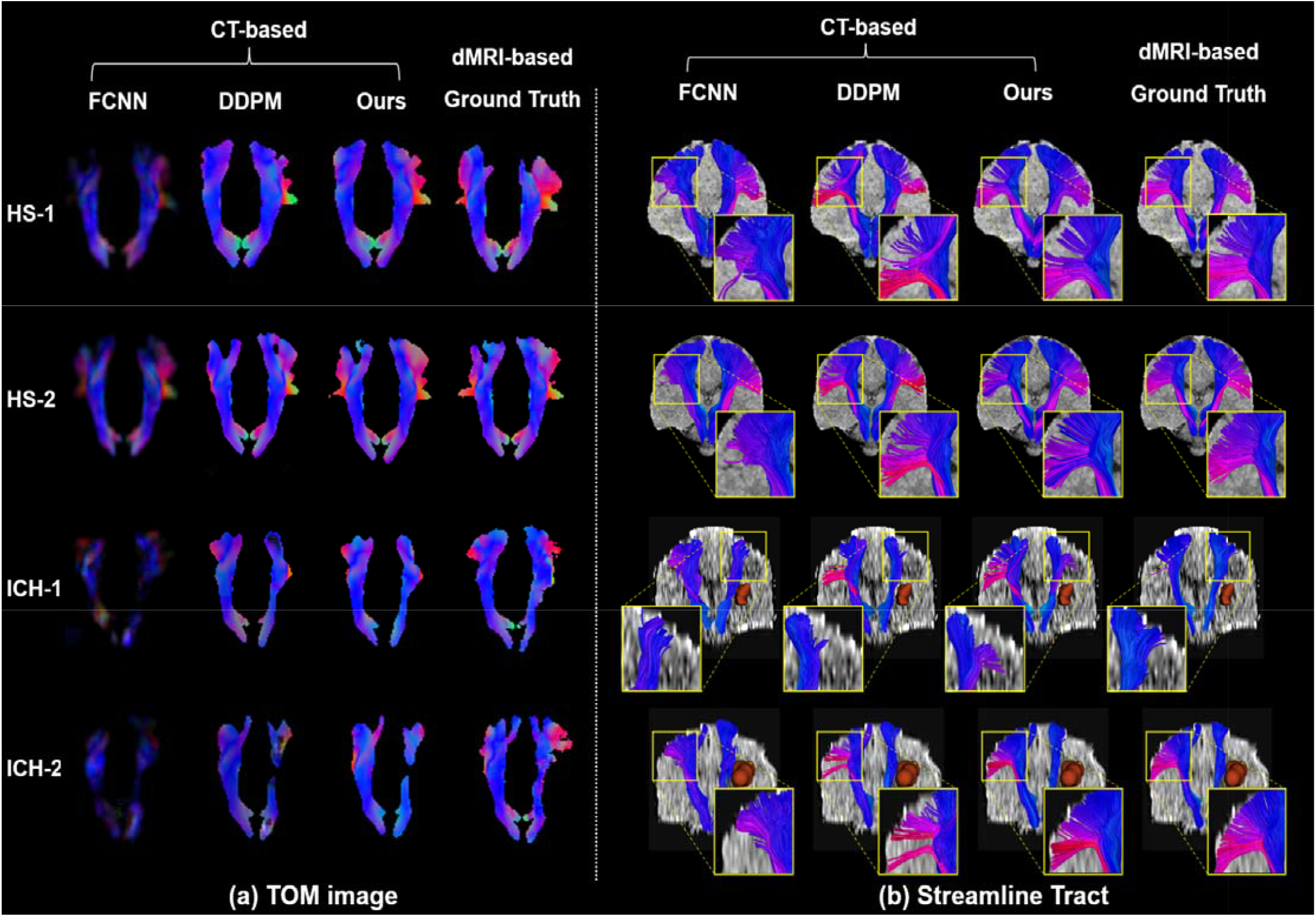
Visual comparison of the predicted tract orientation mapping (TOM) and reconstructed streamline tracts of the corticospinal tract (CST) from baseline methods and our proposed method, with reference to the diffusion MRI (dMRI)–derived ground truth. Results are shown for four example subjects: two healthy subjects (HSs) and two patients with intracerebral hemorrhage (ICH). Baseline methods: FCNN = fully convolutional neural network, DDPM = denoising diffusion probabilistic model. **Note**— In the visualization, yellow rectangles highlight locations with pronounced differences in tract reconstruction. Zoomed insets are provided for detailed inspection of local tract details. In the ICH patients, the red 3D model denotes the ICH location.

### 4.3. Expert Assessment

Figure 3 presents expert ratings of predicted CSTs in the ICH-affected hemispheres based on agreement with two statements: (1) the tract passes through expected brain regions, and (2) ICH-induced CST displacement or splitting is accurately represented. The CT-based method receives consistently favorable evaluations, with high agreement (“strongly agree” or “agree”) rates (statement 1: 61.7%, 37/60; statement 2: 60.0%, 36/60) and low disagreement (“strongly disagree” or “disagree”) rates (both statements: 16.7%, 10/60). Violin plots show significantly higher scores for CT-based predictions compared with dMRI-based results (statement 1: 3.63 vs. 3.25, p=0.017; statement 2: 3.65 vs. 3.30, p=0.028). Inter-expert agreement is high across all assessments (ICC>0.8). Figure 4 shows CT-based versus dMRI-based predictions in example cases, where we observe more complete tract coverage near lesions, whereas the corresponding dMRI-based results are incomplete or absent in these regions.

**Figure 3.**
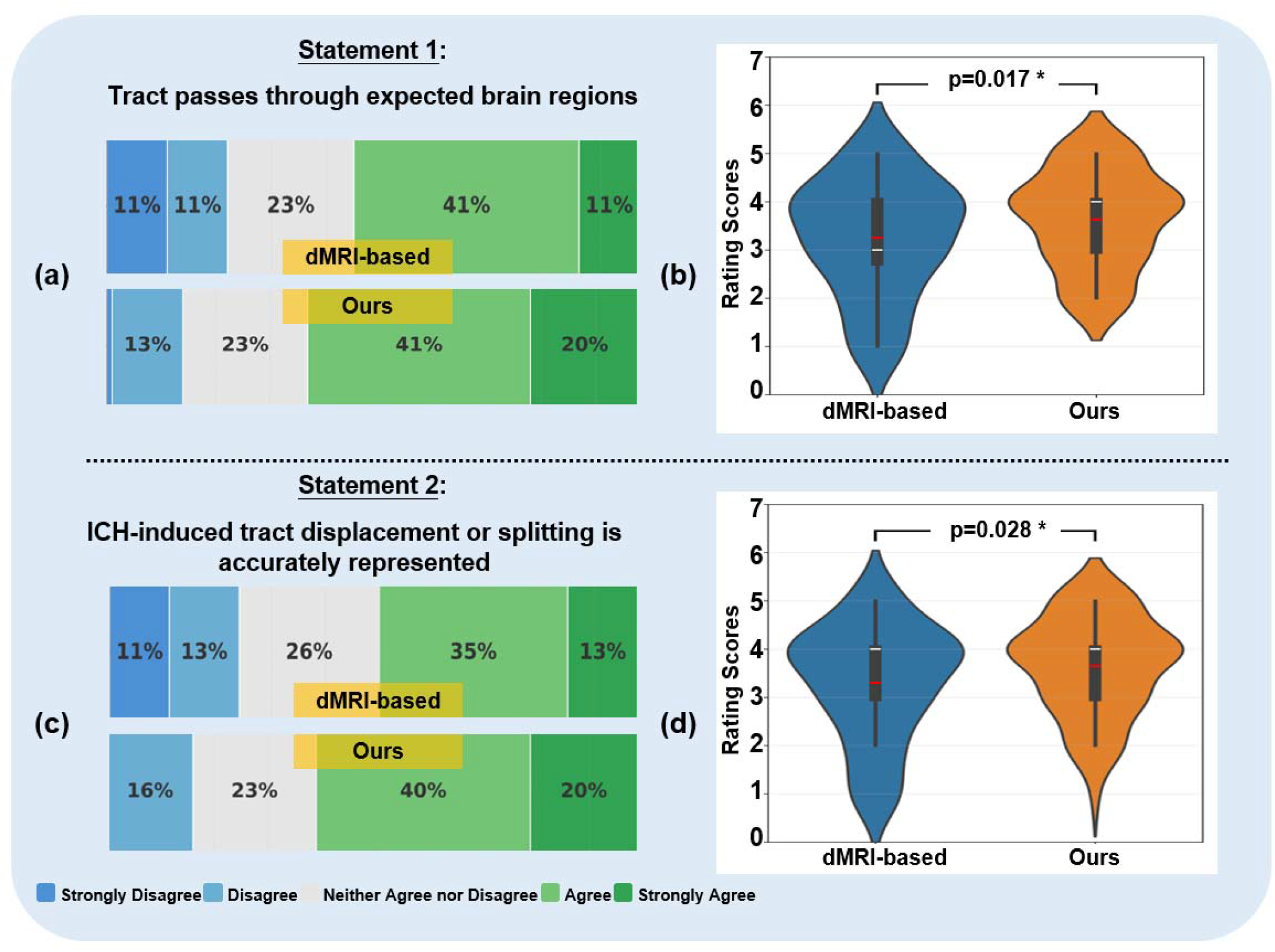
Expert neurosurgeon assessment results comparing the diffusion MRI (dMRI)–based and our proposed CT-based corticospinal tract (CST) reconstruction methods. **(a, c)** Gantt charts display the percentage of experts agreeing with each assessment statement regarding the anatomical plausibility. **(b, d)** Corresponding violin plots illustrate the distributions of the detailed rating scores (1–5 scale, from “Strongly Disagree” to “Strongly Agree”). For statement 1, the mean rating score was 3.63 for CT-based reconstructions, compared to 3.25 for dMRI-based reconstructions. For statement 2, the mean score was 3.65 for CT-based and 3.30 for dMRI-based reconstructions. The CT-based method received significantly higher ratings than the dMRI-based method for both statements (both p < 0.05).

**Figure 4.**
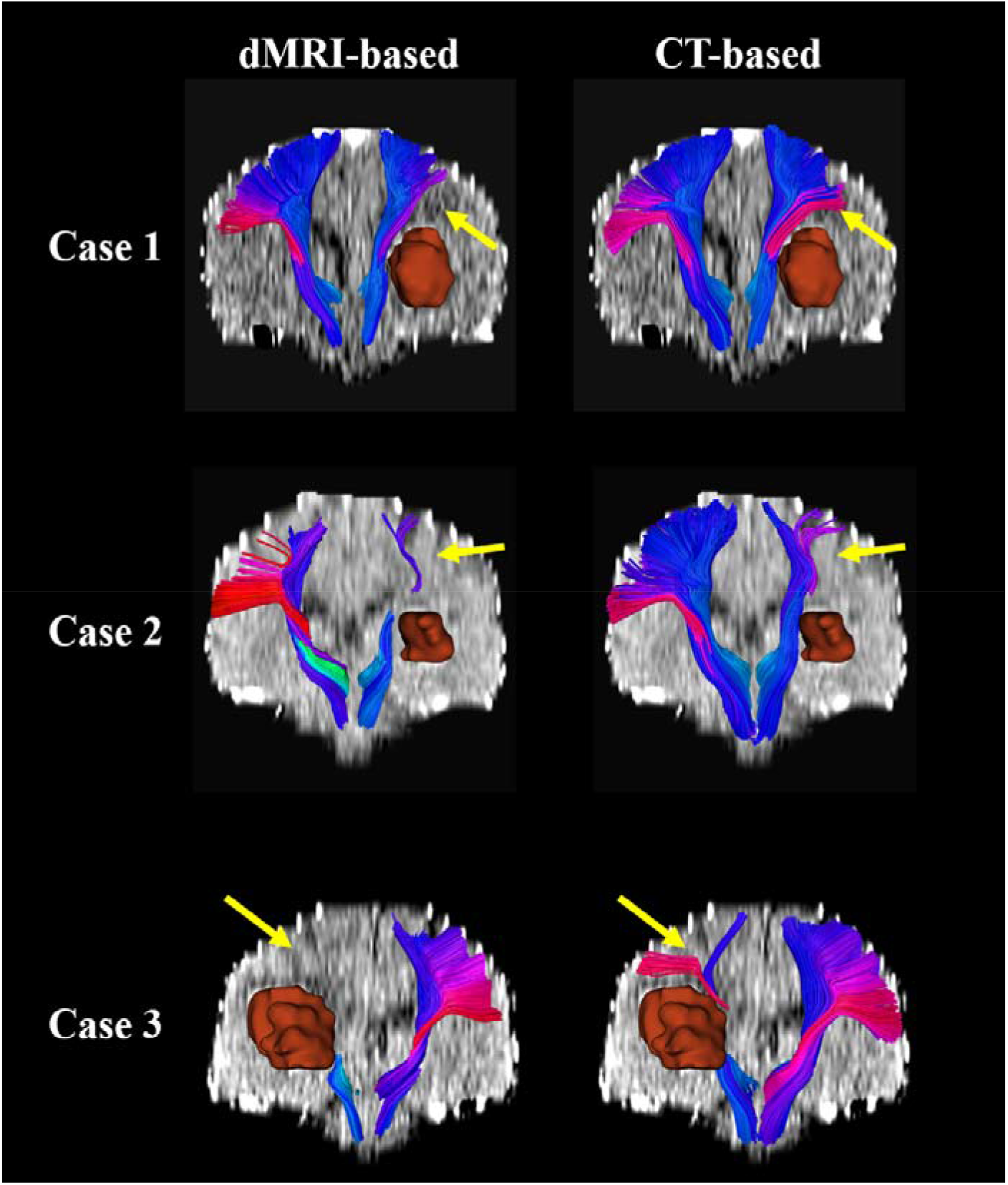
Comparative visualization of corticospinal tract (CST) tractography in three example patients with intracerebral hemorrhage (ICH). For each patient, tractography results from the compared diffusion MRI (dMRI)–based method (left column) are compared with those from the proposed CT-based method (right column). Patient demographic and clinical details: **Case 1**: 47-year-old male with a left basal ganglia hemorrhage (volume, 18 mL); **Case 2**: 52-year-old male with a left basal ganglia hemorrhage (volume, 13 mL); **Case 3**: 46-year-old female with a right basal ganglia hemorrhage (volume, 32 mL). **Note**— In the visualization, the red 3D model denotes the ICH location, and the yellow arrow highlights an example tract location where reconstruction differences between the two methods are visually apparent.

### 4.4. Tract-Behavior Correlation Analysis

Figure 5 shows correlations between muscle strength scores and CST volumes derived from CT- and dMRI-based tracts. Both methods show significant positive correlations, with a stronger association for CT-based CSTs (*r*=0.726, p=3.731×10□ □) than for dMRI-based CSTs (*r*=0.594, p=1.053×10□ □), indicating a closer relationship between CT-based CST integrity and limb motor function in this cohort.

**Figure 5.**
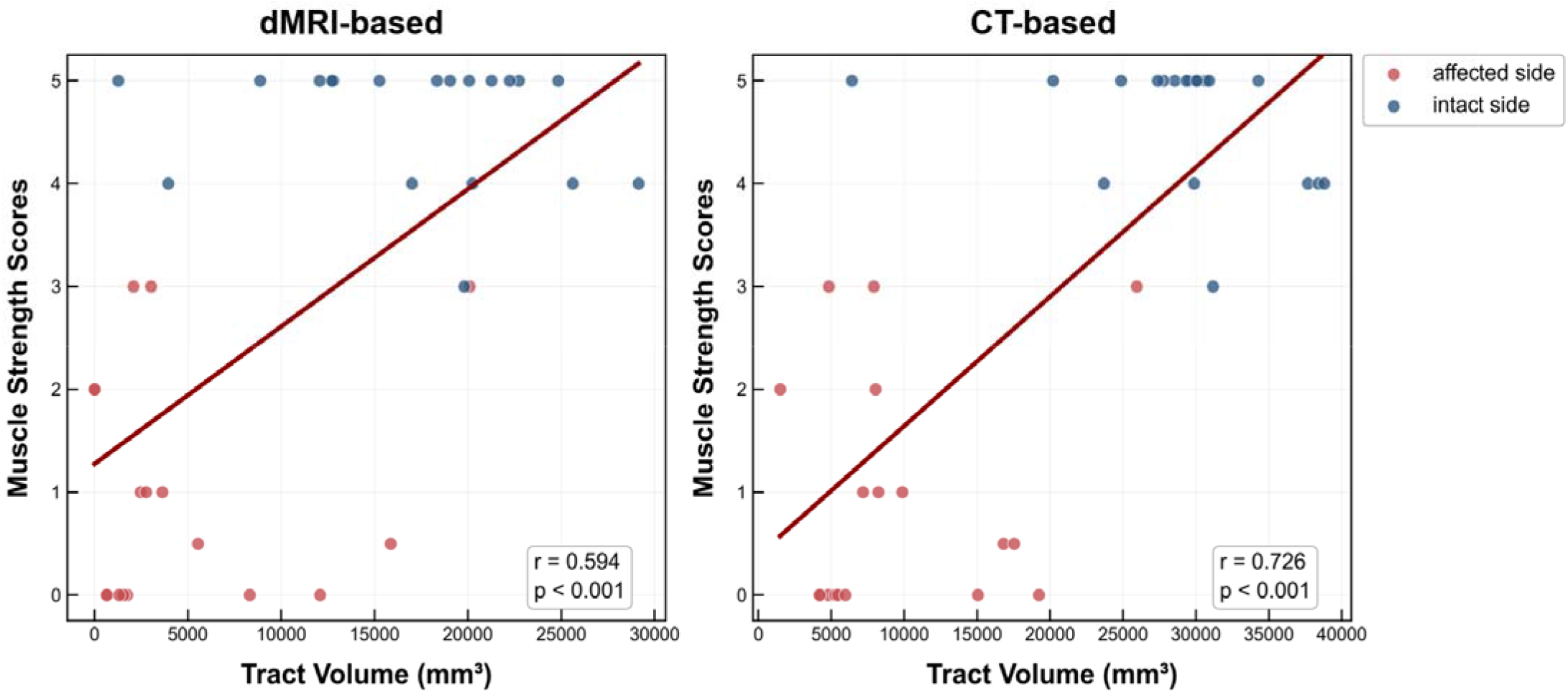
Scatter plots of corticospinal tract (CST) volume versus muscle strength scores, with fitted regression lines. **Left**: Correlation for tract volumes derived from the diffusion MRI (dMRI)-based method. **Right**: Correlation for tract volumes derived from the proposed CT-based method. Both methods demonstrated a statistically significant correlation (dMRI: r=0.594, p<0.001; CT: r=0.726, p<0.001), with a stronger correlation observed for the CT-based method.

## 5. Discussion

This study presents a deep generative model for predicting WM tracts, specifically the CST, directly from routine clinical CT scans in ICH patients. Our approach adopts a TractSeg-inspired strategy: rather than attempting to extract intrinsic structural signals from CT intensities, the model learns to map CT features onto the TOM derived from dMRI. We demonstrate strong tract prediction performance enabled by a segmentation-guided conditioning design that incorporates tract location as an anatomical prior. Our method significantly outperforms baseline architectures (FCNN and DDPM), producing superior tractography both quantitatively and visually. We further show that augmenting ICH patient data with HS data significantly improves performance, likely due to increased sample size and anatomical diversity. Expert assessment demonstrates the reliability of the CT-based CST prediction, as evidenced by its high anatomical plausibility. The correlation analysis indicates a significant correlation between tract integrity and motor function, showing the high clinical relevance of the derived CST.

We demonstrate the superiority of CT-based CST reconstructions over dMRI-based results in the context of ICH. This is evidenced by the higher expert assessment scores and stronger tract-behavior correlations. In patients where the CST is substantially affected by ICH lesions, our CT-based method provides more complete tract coverage in peri-lesional regions than dMRI-based approaches. While the compared TractSeg enables robust dMRI-based tract identification in ICH patients, the presence of hemorrhage and edema can affect the dMRI signals, obscuring the underlying WM tract and impairing tract reconstruction completeness. In contrast, our CT-based approach, which is less sensitive to such signal changes and learns from normal WM tract anatomy, achieved better coverage and consequently higher expert ratings.

Our method demonstrates the viability of a dMRI-free tractography strategy using clinically routine CT. While recent advances have enabled WM tract reconstruction from anatomical T1w data, we extend this paradigm to the more challenging domain of non-contrast CT, specifically in patients with ICH, where lesions critically distort WM structures. Compared to related work by Murray et al. ^25^, which predicts a probabilistic map of CST location from CT, our approach offers two key advances. First, it achieves higher spatial prediction accuracy (Dice: 0.81 vs. 0.57). Second, and more importantly, our method enables full 3D fiber tracking to reconstruct the continuous trajectory of the pathway, moving beyond a probabilistic localization to a tractographic model.

The proposed CT-based approach offers a feasible framework for approximating CST trajectories, presenting a potential alternative in clinical scenarios where dMRI is inaccessible. This method may provide preliminary anatomical insights for patients with acute hematomas in resource-limited settings or for those with MRI contraindications, such as implanted metallic devices or postoperative shunts. By leveraging the widespread availability and rapid acquisition of CT, this technique could offer a more accessible, albeit proxy, measure of tract geography. Furthermore, evaluating estimated CST involvement early after a hemorrhage might provide useful exploratory data regarding motor recovery, potentially aiding in clinical discussions and the refinement of rehabilitation strategies. While it does not replace the gold-standard precision of dMRI, this CT-based mapping serves as a pragmatic tool for enhancing our understanding of white matter disruption when advanced imaging is not an option.

This study has several limitations that point to future research directions. First, because of the difficulty in acquiring paired CT-dMRI datasets, our cohort was limited to a single-center population. Future work could validate the generalization of our method to multi-center data. Second, while we focused on the CST due to its clinical relevance in ICH surgery, future investigations could extend this methodology to other WM tracts affected by ICH. Third, further optimization of the network architecture is warranted to improve tract prediction performance.

In conclusion, we have developed a novel learning-based approach that enables direct tractography from CT scans. This provides an efficient and practical solution for preoperative WM mapping in ICH, offering the key advantages of tractography in resource-limited or time-critical clinical settings.

## Acknowledgements

This work is in part supported by the National Natural Science Foundation of China (No. 62371107) and the National Key R&D Program of China (No. 2023YFE0118600).

## CRediT authorship contribution statement

**Guanlin Huang:** Writing - review & editing, Writing - original draft, Visualization, Validation, Methodology, Investigation, Formal analysis, Data curation; **Guoqiang Xie:** Writing - review & editing, Data curation, Resources, Validation; **Yijie Li:** Writing - review & editing; **Qun Wang:** Writing - review & editing, Validation; **Shun Yao:** Writing - review & editing, Validation; **Yiheng Tan:** Writing - review & editing, Validation; **Ron Kikinis:** Writing - review & editing, Software; **Alexandra J Golby:** Writing - review & editing; **Lauren J O’Donnell:** Writing - review & editing; **Fan Zhang:** Writing - review & editing, Writing - original draft, Resources, Methodology, Investigation, Funding acquisition, Formal analysis, Data curation.

## Data availability

The datasets are not publicly available because of hospital regulations and patient privacy but are available from the corresponding author on reasonable request.

## Code Availability

The source code is available at GitHub (https://github.com/PGHGlin/White-Matter-Tract-Mapping-in-Intracerebral-Hemorrhage).

